# A bacterial sialidase mediates early life colonization by a pioneering gut commensal

**DOI:** 10.1101/2023.08.08.552477

**Authors:** Ekaterina Buzun, Chia-Yun Hsu, Kristija Sejane, Renee E. Oles, Adriana Vasquez Ayala, Luke R. Loomis, Jiaqi Zhao, Leigh-Ana Rossitto, Dominic McGrosso, David J. Gonzalez, Lars Bode, Hiutung Chu

## Abstract

The early microbial colonization of the gastrointestinal tract can lead to long-term impacts in development and overall human health. Keystone species, including *Bacteroides spp*., play a crucial role in maintaining the structure, diversity, and function of the intestinal ecosystem. However, the process by which a defined and resilient community is curated and maintained during early life remains inadequately understood. Here, we show that a single sialidase, NanH, in *Bacteroides fragilis* mediates stable occupancy of the intestinal mucosa and regulates the commensal colonization program during the first weeks of life. This program is triggered by sialylated glycans, including those found in human milk oligosaccharides and intestinal mucus. After examining the dynamics between pioneer gut *Bacteroides* species in the murine gut, we discovered that NanH enables vertical transmission from dams to pups and promotes *B. fragilis* dominance during early life. Furthermore, we demonstrate that NanH facilitates commensal resilience and recovery after antibiotic treatment in a defined microbial community. Collectively, our study reveals a co-evolutionary mechanism between the host and the microbiota mediated through host-derived glycans to promote stable intestinal colonization.

## INTRODUCTION

Microbial colonization of the infant gut has important impacts on childhood development and overall human health. This early life colonization event can resolve into long-lasting consequences, modulating the risk of allergies, asthma, and metabolic disorders^1^. While multiple factors shape the dynamics and composition of the developing microbial community, the infant microbiota is strongly influenced by the delivery mode^2–4^ and diet^5,6^. Numerous studies have established that consumption of human milk influences the microbial community of the infant gut, promoting the growth of bacteria (e.g., *Bifidobacterium* and *Bacteroides*) capable of degrading human milk oligosaccharides (HMOs). While the relationship between *Bifidobacterium* and HMOs has been the primary focus of investigation^7–9^, there is now mounting evidence that *Bacteroides* species serve as pioneer species of the infant microbiome within the first days of life^10–12^.

*Bacteroides* species possess an extraordinary repertoire of polysaccharide utilization loci (PULs) that facilitate the breakdown of complex host-derived glycans, including mucins and HMOs^13–15^. Studies with *Bacteroides thetaiotaomicron* identified PULs that mediate foraging of intestinal mucus to support colonization and persistence in the gut^15,16^. Moreover, previous work with *Bacteroides fragilis* demonstrated the commensal colonization factor (CCF) locus enables long-term colonization of the murine gut^17,18^. The CCF system that mediates *B. fragilis* occupancy in the mucosal niche is classified as PUL^17^, suggesting a role of glycan utilization in directing intestinal colonization. In contrast to commensal colonization, our understanding of how pathogenic bacteria leverage endogenous host glycans to colonize and invade is considerably more comprehensive^19^. Bacterial sialidases, for instance, contribute to colonization of mucosal sites by providing nutrients through sialic acid catabolism, unmasking docking sites for adherence and biofilm formation^20,21^. However, less is known with regards to whether host glycans, such as HMOs, directly promote colonization of commensal bacteria. The ability to metabolize and utilize HMOs presumably would provide an advantage for these commensals to stably colonize the infant gut. Several studies have shown that *Bacteroides* are capable of degrading HMOs^22,23^, however, the mechanisms directly linking HMO metabolism to commensal colonization of the infant gut remain undetermined.

Infants are rapidly colonized with microbes during birth and soon after begin to consume a diet of human milk and/or infant formula^5,10^, exposing the environment of the infant gut to HMOs. Importantly, HMOs share structural similarities with intestinal mucus O-glycans^24^. This supports a model where HMOs act as the primary signal for pioneer commensals to initiate gut colonization during early life, after which mucosal glycans serve to maintain persistence and resilience throughout life. Here, we define the HMO utilization profile in *B. fragilis* and reveal its selective preference for sialylated HMOs. Through comparative proteomics, we unveil the metabolic system governing HMO utilization in *B. fragilis* and identify the induction of a sialidase (NanH) and the CCF locus, suggesting HMO metabolism and intestinal niche occupancy are tightly linked. Our findings demonstrate that the *B. fragilis* sialidase, NanH, is crucial for stable colonization in the murine gut, as isogenic mutants are outcompeted by wildtype (WT) *B. fragilis* in the colonic lumen and mucosal compartments. Moreover, *B. fragilis* abundance in pups during the nursing period is dependent on NanH expression. Notably, the role of NanH in colonization extends beyond suckling pups, as mutants lacking NanH sialidase exhibit impaired competitive fitness during co-colonization with *B. thetaiotaomicron* in adult mice. Finally, we demonstrate that in a complex community, *B. fragilis* can persist and rebound following antibiotic treatment in a NanH-dependent manner. Our results provide the first evidence that host-derived glycans, such as HMOs, act as the primary signal for pioneer commensals to initiate gut colonization during early life, facilitating their persistence in the intestinal mucosa throughout life.

## RESULTS

### Defining the HMO metabolizing system in *Bacteroides fragilis*

We first assessed the ability of *Bacteroides* species to utilize HMOs as a sole carbon source. Using pooled HMOs (pHMOs) purified from donor human milk, the growth of *B. fragilis* (NCTC 9343), *B. thetaiotaomicron* (VPI-8482), *B. vulgatus* (ATCC 8482), *B. ovatus* (ATCC 8483), *B. salyersie* (DSM 18765), *B. uniformis* (ATCC 8492), and *B. acidifaciens* (JCM 10556) was monitored continuously for 72 hours. From these seven *Bacteroides* species, *B. fragilis*, *B. thetaiotaomicron*, and *B. vulgatus* displayed robust growth, whereas *B. ovatus*, *B. salyersie*, *B. uniformis*, and *B. acidifaciens* exhibited slow growth (Figure 1A). Thin layer chromatography (TLC) analysis revealed that the composition of products remaining from stationary phase cultures varied among *Bacteroides* species (Figure 1B). In contrast to *B. thetaiotaomicron* and *B. vulgatus*, *B. fragilis* was the most efficient and displayed a distinct ability to rapidly deplete available HMOs (Figure 1B). To further dissect the HMO utilization profile in *B. fragilis* we used high performance liquid chromatography (HPLC) to separate and identify 19 most abundant HMO structures (Figure S1A). *B. fragilis* was grown on 15 mg/ml pHMOs and the HMO utilization profile was assessed by HPLC (Figure 1C) and TLC (Figure S1B) at regular intervals for 48 hours. This revealed that *B. fragilis* prioritized consumption of ɑ2-6-sialylated HMOs, including 6’-sialyllactose (6’SL), disialyllacto-N-tetraose (DSLNT), and disialyllacto-N-hexaose (DSLNH) (Figure 1C). Altogether, these data suggest that *B. fragili*s possesses a coordinated metabolic system, demonstrating a selective preference for sialylated HMOs.

**Figure 1.**
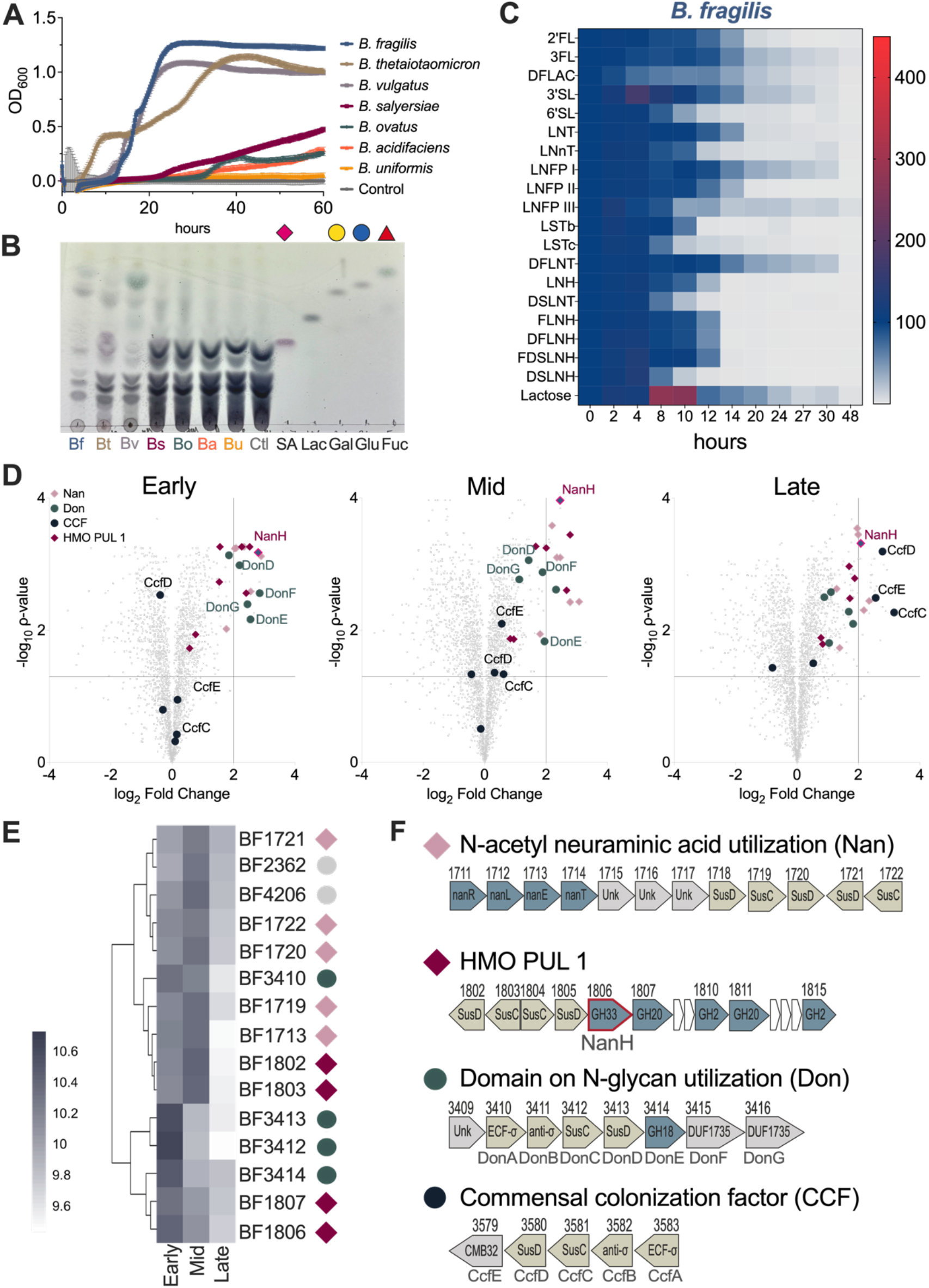
Defining HMO utilization system in *B. fragilis*. (A) Eight *Bacteroides* species *B. fragilis* NCTC 9343 (Bf), *B. thetaiotaomicron* VPI-8482 (Bt), *B. vulgatus* ATCC 8482 (Bv), *B. ovatus* ATCC 8483 (Bo), *B. salyersie* DSM 18765 (Bs), *B. uniformis* ATCC 8492 (Bu), and *B. acidifaciens* JCM 10556 (Ba) were grown in defined medium supplemented with 1.5 % pooled HMOs (pHMOs). (B) TLC of cell-free supernatant collected from stationary phase cultures of eight *Bacteroides* species on pHMOs. Controls include SA – sialic acid/Neu5AC (1 μM); Lac – lactose (1 mg/ml); Gal – galactose (1 μM), Glc – glucose (1 μM); Fuc – fucose (1 μM). (C) Selective degradation of pHMOs by *B. fragilis* over time. Cell free supernatant was analyzed by HPLC, at each timepoint peaks were normalized to the internal control and compared to abundance at t=0. (D) Proteomic analysis of pHMO-grown whole *B. fragilis* cells, mass spectral data was normalized to glucose-grown cells harvested at mid-log phase. pHMOs vs glucose fold change at early-, mid-, and late-log phases. Significance was based on Log_2_ FC ≥2, *P*-value < 0.05. Colors and shapes correspond to proteins grouped in abundant operons. (E) Normalized relative abundance of proteins with a max log_10_ fold change ≥2 and a Benjamini Hochenburg corrected *P*-value < 0.05 between samples across growth phases. Hierarchical clustering was performed using Euclidean distance. Scale bar shows protein abundance from high (purple) to low (white). (F) Organization of significantly abundant polysaccharide utilization loci (PULs) in *B. fragilis* during growth on pHMO, gene labeling corresponds to original locus tags BFxxxx: N-acetyl neuraminic acid (sialic acid) utilization (Nan) operon (pink diamond); HMO PUL1 (maroon diamond); Domain of N-glycan (Don) utilization operon (teal circle); Commensal colonization factor (CCF) operon (black circle).

We then sought to uncover the metabolic system that enables selective HMO metabolism in *B. fragilis*. Importantly, *B. fragilis* displayed a biphasic growth on pHMOs (Figure 1A), during which bacteria are known to shift their metabolism as a means of adaptation to environmental changes^25–27^. TLC and HPLC analyses of the culture supernatants from the early-, mid-, and late-log growth phases showed that indeed the substrate diversity and composition gradually changed over time (Figures S1C-E). To identify the *B. fragilis* protein repertoire orchestrating HMO metabolism during biphasic growth, we performed whole cell proteomics on HMO-grown *B. fragilis* at the three growth phases. In total, 2,808 proteins were identified (Figure 1D, Figure S2A, and Table 1). The number of significantly abundant proteins (>2-fold change compared to glucose control, *P*<0.05,) varied across the three time points, demonstrating a metabolic shift in *B. fragilis* during consumption of HMOs (Figure 1D). Notably, a group of proteins was consistently abundant in *B. fragilis* over all three growth phases (Figure 1D and 1E). This included components of PUL 30^28^ (hereafter HMO PUL 1) which encodes a sialidase BF1806 (NanH, GH33); two β-N-acetylglucosaminidase, BF1807 and BF1811 (GH20); one β-galactosidase, BF1815 (GH2); one putative β-mannosidase, BF1810 (GH2); and two pairs of SusC and SusD proteins (Figure 1D-F). Additionally, the Nan operon (PUL 20), which mediates the uptake of Neu5Ac (sialic acid) cleaved by the sialidase NanH and its conversion into glucose-6-phosphate for downstream glycolysis^29,30^, was also consistently abundant (Figure 1D-F). We further validated the upregulated expression of the HMO PUL 1 genes, *bf1802* (SusD), *bf1806* (*nanH*, GH33), and *bf1807* (GH20), in *B. fragilis* during growth on pHMOs (Figure S2B). These data suggest that the HMO PUL 1 is the central enzymatic system mediating HMO breakdown in *B. fragilis*.

Further analysis demonstrated that proteins comprising the N-glycan utilization (Don) operon^31^ (PUL 43)^28^ resided in the top 0.5% of the total protein repertoire in the early- and mid-log phases of growth (Figure 1D-F). During the metabolic shift associated with the late-log phase growth, the Don operon was displaced with proteins involved in intestinal colonization, including the commensal colonization factor (CCF) locus^17^ (Figure 1D-F) and the fucRIOAKP operon^32^ (Figure S2C). Both the Don and CCF operons enclose putative proteins, BF3415 (DonF), BF3416 (DonG), and BF3579 (CcfE), sharing a domain of unknown function DUF1735, which is annotated as a carbohydrate binding module (CBM) 32 in PUL-DB^28^. Further, all three proteins are predicted to be cell surface associated and thus most likely mediate glycan binding and acquisition. Interestingly, out of the 100 most upregulated proteins in the late-log phase, 40% accounted for lone SusC-SusD pairs and proteins with domains of unknown function (DUFs) predicted to act as glycan binding modules (Figure S2D). We therefore propose *B. fragilis* uses a range of binding proteins to capture extracellular HMOs and perform their metabolism intracellularly, while using the sialidase NanH (BF1806) and other enzymes in HMO PUL 1 as a central enzymatic system for depolymerization of HMOs.

### NanH sialidase promotes intestinal colonization in a commensal bacterium

Our work here (Figure 1) demonstrates that *B. fragilis* upregulates a sialidase, NanH (BF1806), in response to pHMOs. Phylogenetic analysis indicated that homologues of BF1806 can be found in at least 11 *Bacteroides* species and a number of other commensals (Figure S3A). However, these enzymes are enclosed in distinct operons, sharing low similarity with NanH (BF1806), with the closest homologues present in *B. intestinalis, B. salyersiae*, and *B. nordii* (Figure S3B and S3C). To determine the functional role of *B. fragilis* NanH in HMO metabolism, we generated an isogenic *B. fragilis* mutant lacking BF1806 (Δ*nanH*). No differences in growth were observed in nutrient-rich media (BHI-S) between *B. fragilis* WT and the Δ*nanH* mutant (Figure 2A). We next assessed the ability of Δ*nanH* to grow on pHMOs and revealed delayed growth on pHMOs (WT AUC=27 vs Δ*nanH* AUC=13.13, *P*<0.001) (Figure 2B). Notably, 20% of pHMOs contain ɑ2-3/6-Neu5AC decorations and thus can be separated into sialylated (acidic) and non-sialylated (neutral) HMOs (Figure 2C). We then fractionated pooled human milk samples and demonstrated that *B. fragilis* Δ*nanH* strain, lacking its central sialidase, failed to utilize and grow in pooled sialylated HMOs (WT AUC=13.93 vs Δ*nanH* AUC=7.6, *P*<0.01) (Figure 2D), while its ability to use pooled non-sialylated/neutral HMOs was not attenuated (WT AUC = 8.5 vs Δ*nanH* AUC = 7.9) (Figure 2E). Consistently, HPLC analyses of the supernatant after growth revealed that Δ*nanH* was unable to utilize 6’SL and exhibited partial degradation of other HMOs in the heterogeneous pHMOs sample (Figure S4A and S4B), further corroborating that NanH is required for the metabolism of sialylated HMOs in the heterogeneous pool. To investigate linkage specificity of the NanH sialidase, we grew *B. fragilis* WT and Δ*nanH* in singular HMOs, 3’SL and 6’SL, harboring ɑ2-3- and ɑ2-6-Neu5AC decorations, respectively. This further demonstrated that the Δ*nanH* mutant was unable to grow on 6’SL (WT AUC=3.8 vs Δ*nanH* AUC not detected) (Figure 2F) and exhibited significantly impaired growth kinetics on 3’SL (WT AUC=5.7 vs Δ*nanH* AUC=2.1, *P*<0.001; Figure 2G). Combined, our findings demonstrate that *B. fragilis* NanH is a central sialidase and displays activity against ɑ2-6/ɑ2-3-Neu5Ac decorations.

**Figure 2.**
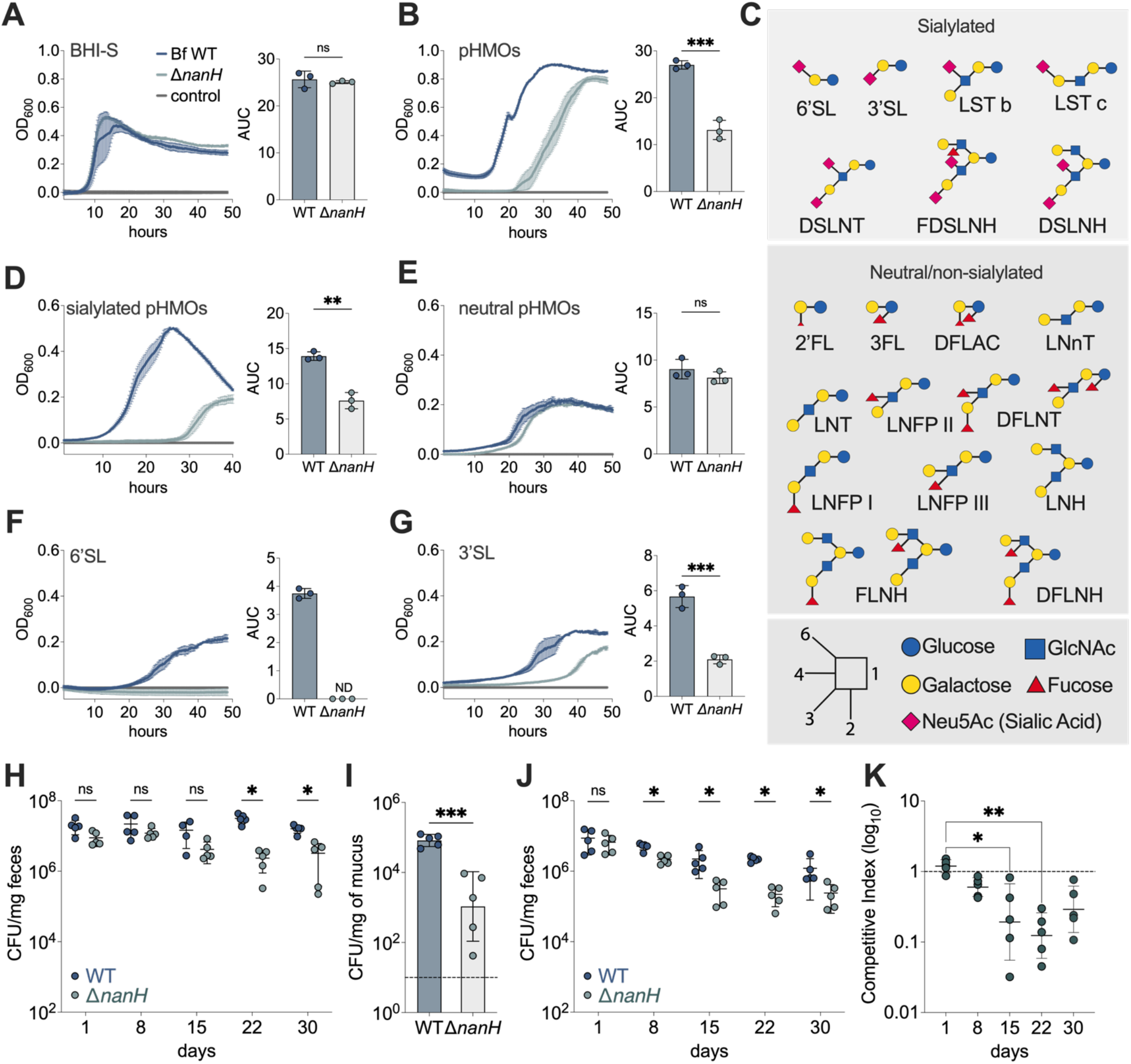
NanH is required for growth on HMOs and promotes intestinal colonization. (A – G) Growth curves and associated area under the curve (AUC) of *B. fragilis* WT (dark blue) and Δ*nanH* isogenic mutant (light blue) on (A) BHI-S; (B) pooled HMOs (pHMOs); (C) Diagrammatic structures of Neutral/non-sialylated (top) and sialylated (bottom) HMOs. Sugars are color coded as indicated, the linkage schematics is shown in the bottom left corner; (D) pooled sialylated HMOs; (E) pooled neutral HMOs; (F) 6’-sialylactose (6’SL); (G) 3’-sialylactose (3’SL). Growth curves show mean in solid line ± SD of three biological replicates, representative from two independent experiments. Growth rates were calculated from three biological replicates ± SD (H – I) Germ-free mice were orally gavaged with a single dose of either *B. fragilis* WT or Δ*nanH* at 10E7 CFUs per mouse. (G) Quantification of *B. fragilis* WT and Δ*nanH* in feces of mono-colonized mice over 30 days. (I) Abundance of WT and Δ*nanH* strains in the colonic mucosa 30 days after colonization. The dashed line shows the limit of detection (LOD) 10 CFU/mg (J– K) Germ-free mice were orally gavaged with 1:1 mixture of *B. fragilis* WT and Δ*nanH* strains at 10E7 CFUs of each strain. (J) Quantification of *B. fragilis* WT and Δ*nanH* in feces of co-colonized mice over 30 days. (K) The competitive index (log10) for co-colonization experiments as indicated by input/output ratios. Differentiation between the WT and the isogenic Δ*nanH* mutant was performed using resistance to tetracycline and chloramphenicol conferred by pFD340 plasmids, respectively. *n* = 5, data are representative of at least two independent experiments. Data are shown as geometric mean ± SD. \**P* < 0.05; \*\**P* < 0.01; \*\*\**P* < 0.001; ns – not significant; (Kruskal-Wallis and Mann-Whitney tests); ND – not detected.

Importantly, gastrointestinal mucin glycans are abundantly sialylated and the extent of sialylation varies throughout anatomical sites^24,33^. Moreover, sialylated HMOs share structural similarities with glycans decorating mucus glycoproteins^24^ and previous studies reported that *B. fragilis* NanH is highly expressed during growth on porcine gastric mucin (PGM) II^16^. Building on our finding that Δ*nanH* mutants are deficient in utilizing sialylated host glycans (Figure 2D-G) and the induction of the CCF locus during growth on pHMOs (Figure 1D-F), we investigated whether the NanH sialidase may also play a role in niche occupancy within the intestinal mucosa. Through mono-colonization of germ-free (GF) C57BL/6J mice, we observed a steady decrease in bacterial numbers of the Δ*nanH* isogenic mutant over time, failing to establish full colonization when compared to *B. fragilis* WT (∼10-fold decrease; *P*<0.001) (Figure 2H). Moreover, the abundance of the Δ*nanH* mutant was significantly reduced in the colonic mucus in comparison to *B. fragilis* WT mono-colonized mice (∼100-fold reduction, *P*<0.001) (Figure 2I), suggesting that NanH sialidase indeed mediates mucosal occupancy in the gut. To assess the competitive fitness of *B. fragilis* Δ*nanH*, we orally gavaged germ-free mice with *B. fragilis* WT and the isogenic Δ*nanH* mutant at 1:1 ratio. *B. fragilis* WT displayed a significant competitive advantage over Δ*nanH* at 15 days post-co-colonization (∼100-fold decrease, *P*<0.001) (Figure 2J and 2K), indicating that the presence of a functional NanH contributes to commensal fitness *in vivo*.

Our proteomics data indicated that pHMOs induce expression of the commensal colonization factor, CCF (Figure 1), which was previously shown to facilitate occupancy of the intestinal crypts^17^. We next investigated if the inability of the Δ*nanH* strain to colonize the intestinal mucosa is linked to the CCF locus and analyzed relative expression of the key *ccf* genes: *ccfC*, *ccfD*, and *ccfE*. qPCR analysis of the fecal content and colonic mucosa collected 21 days after mono-association with the Δ*nanH* mutant revealed that the relative expression of *ccfC*, *ccfD*, and *ccfE* was significantly decreased compared to the *B. fragilis* WT in mono-colonized mice (Figure S5A and 5B). Notably, the expression of *ccfC* and *ccfD* was reduced in Δ*nanH* during growth on pHMOs *in vitro* (Figure S5C). These data suggest a functional link between the NanH sialidase and the expression of genes comprising the CCF locus, further supporting the role of NanH in commensal colonization. Although the absence of the CCF does not impair the ability of *B. fragilis* to utilize pHMOs, the double mutant Δ*nanH*ΔCCF exhibited a more pronounced growth defect on pHMOs than Δ*nanH* single mutant (Figure S5D and 5E). This finding indicates that NanH and the CCF locus work synergistically during pHMO metabolism, with NanH acting upstream. Combined, these data suggest that the NanH sialidase orchestrates metabolism of sialylated host glycans, functions as a colonization factor, and governs the activation of the commensal colonization program in *B. fragilis*.

### *B. fragilis* NanH sialidase determines stable niche occupancy and colonization during early life

Previous work demonstrated *B. fragilis* can stably occupy its niche within the intestinal mucosa and concurrently permit colonization by other gut *Bacteroides* (e.g., *B. thetaiotaomicron* or *B. vulgatus*), but not isogenic *B. fragilis* strains^17^. Access to the nutritional mucosal niche requires the removal of the terminal sialic acid moieties that cap the colonic mucus glycans. We thus explored whether *B. fragilis* requires the sialidase NanH to achieve stable niche occupancy of the intestinal mucosa. Germ-free mice were either first mono-associated with *B. fragilis* WT, followed by a challenge with Δ*nanH* on day 8 (Figure 3A), or first mono-associated with Δ*nanH* followed by *B. fragilis* WT challenge (Figure 3B). Indeed, when *B. fragilis* WT served as the initial strain to colonize gnotobiotic mice, the WT strain did not allow for the sequential co-colonization by Δ*nanH* and gradually cleared the mutant strain (Figure 3A). In contrast, Δ*nanH* strain lacked colonization resistance and was permissive to the challenge by *B. fragilis* WT (Figure 3B). Notably, enumeration of the WT and Δ*nanH* strains 25 days post-challenge revealed that Δ*nanH* was present at significantly lower numbers in the feces (Figure 3C) and was undetectable in the mucus of mice that were first colonized with the WT *B. fragilis* (Figure 3D). In contrast, both Δ*nanH* and WT were present in the feces and the mucus of mice that were initially mono-colonized with Δ*nanH* (Figure 3C and 3D), indicating that a functional NanH sialidase is required for the stable occupancy of the intestinal mucosa.

**Figure 3.**
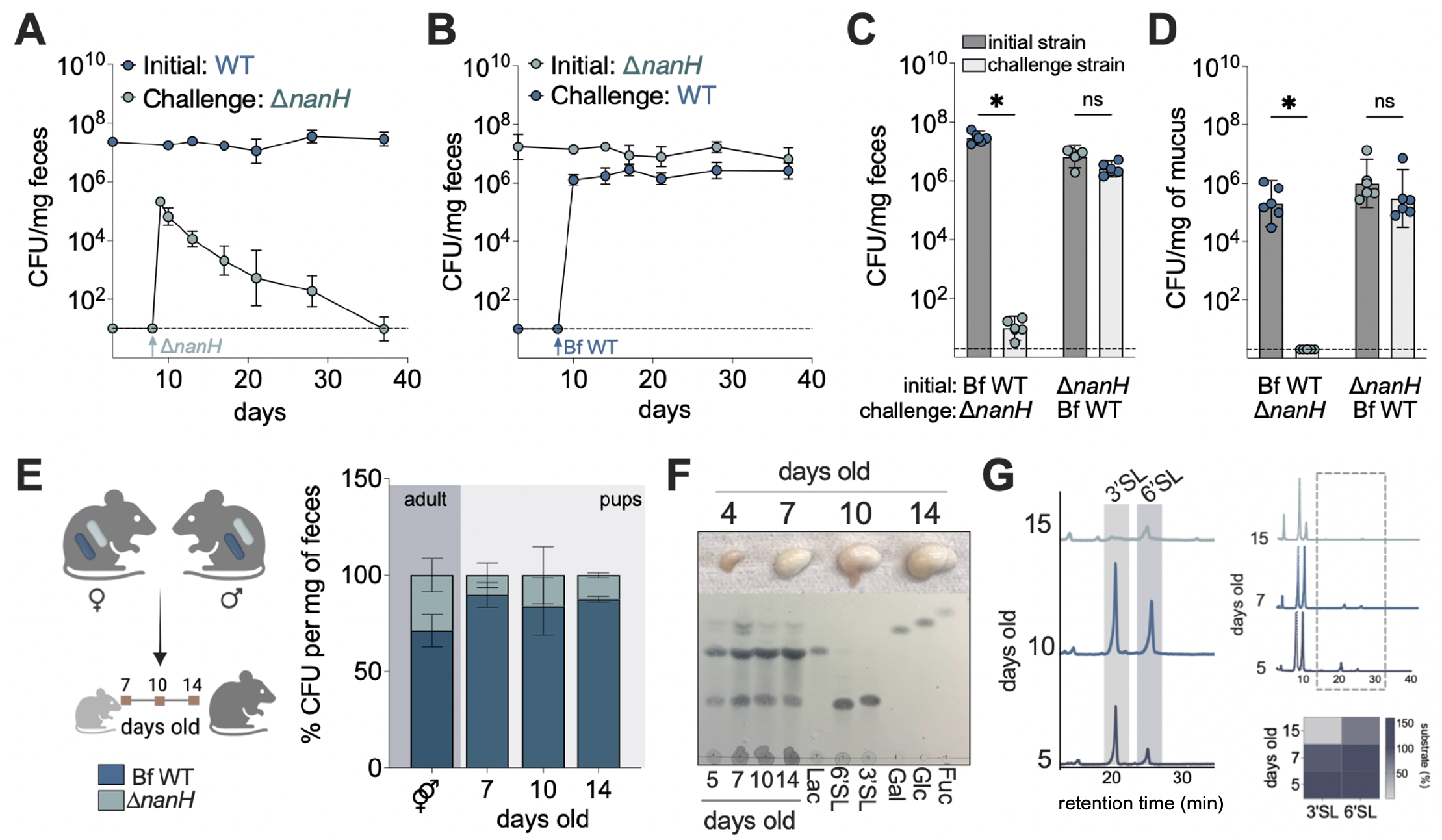
NanH mediates stable niche occupancy and early life colonization. (A) Germ-free mice were mono-associated with *B. fragilis* WT (initial strain) and subsequently challenged with Δ*nanH* strain on day 8. Colony forming units (CFUs) per mg of feces were determined up to day 38 post-colonization. (B) Germ-free mice were mono-associated with Δ*nanH* (initial strain) and challenged with *B. fragilis* WT on day 8. CFUs/mg of feces were determined up to day 38 post-colonization. (C) CFU/mg of the initial and the challenge strains recovered from the fecal pellets on day 27 post-schallenge. (D) CFU/mg of the initial and the challenge strains recovered from the colonic mucus 27 days post challenge. Differentiation between the WT and the isogenic Δ*nanH* mutant was performed using resistance to tetracycline and chloramphenicol conferred by pFD340 plasmids, respectively. *n* = 5, data are representative of at least two independent experiments. Data are shown as geometric mean ± SD. Limit of detection (LOD) is shown in a dashed line, LOD =10-5 CFU/mg; \**P* < 0.05; \*\**P* < 0.01; \*\*\**P* < 0.001; ns – not significant; (Kruskal-Wallis and Mann-Whitney tests); ND – not detected. (E) Vertical transmission of WT and Δ*nanH* from the dam to pups. Abundance of WT and Δ*nanH B. fragilis* in the feces of adult breeders (dark grey shaded area) and the feces of the newborn suckling pups (light grey shaded area). *n* ≧ 3 pups per time point. (F) Top panel: pictures of the stomachs collected from the pups on indicated days. Bottom panel: TLC analysis of the homogenized stomachs resuspended in PBS as indicated. Controls include: 6’SL, 3’SL, lactose (Lac), glucose (Glc), galactose (Gal), and GlcNAc. Trisaccharides and monosaccharides – 1 mg/ml and 1 mM, respectively. (G) HPLC analysis of the stomach contents at indicated time points. Peaks in the shaded area correspond to 3’SL and 6’SL. The heatmap shows % abundance of 3’SL and 6’SL in the stomachs from day 5 (dark purple 150-100%) to day 15 (light gray 2%).

In contrast to HMOs that typically comprise of heterogeneous structures, mouse milk is primarily composed of lactose and sialylated glycans, 3’SL and 6’SL^34,35^, and their abundance gradually declines 14 days postpartum^34^. From our *in vitro* growth data, demonstrating that the lack of NanH resulted in the impaired ability of *B. fragilis* to utilize sialylated HMOs (Figure 2D, 2F, and 2G), we next employed a mouse model to explore whether NanH facilitates *B. fragilis* colonization during the suckling period. Germ-free breeder pairs were orally gavaged with *B. fragilis* WT and Δ*nanH* at equal ratio and upon delivery, newborn pups were sacrificed at 7, 10, and 14 days of age (Figure 3E). *B. fragilis* WT efficiently colonized colons of the newborn pups, dominating over the Δ*nanH* strain, despite both strains being present in the adult breeders at 70% to 30%, respectively (Figure 3E). Further, TLC and HPLC analyses of the stomach contents supported the evidence that the pups received a diet primarily composed of 3’SL, 6’SL, and lactose during the first 14 days of life (Figure 3F and 3G). These data link the ability to utilize sialylated HMOs to the commensal competitive fitness in the intestinal mucosa during early life.

### NanH mediates resilience and competitive fitness of *B. fragilis*

A number commensal bacteria possess sialidases (Figure S3), granting them access to host-derived glycans^24^. However, the specific role of sialidases during competitive co-colonization has not been extensively studied *in vivo*. Notably, *B. thetiotaomicron* expresses a homologue of NanH (BT4055)^22,36^ (Figure S3), which can cleave sialic acids but lacks the *Bacteroides*-specific nanLET operon essential for downstream metabolism of sialic acid^30^. As *Bacteroides* are known to cooperate in the breakdown of complex carbohydrates^13,37^, we assessed the ability of *B. fragilis* WT and Δ*nanH* to share the intestinal niche with *B. thetaiotaomicron* (Bt) during early life. Germ-free breeder pairs were co-colonized with *B. fragilis* WT and *B. thetaiotaomicron* (Bf WT:Bt) or *B. fragilis* Δ*nanH* and *B. thetaiotaomicron* (Δ*nanH*:Bt) for 2-3 weeks and the pups were sacrificed on days 7, 9, and 14 to determine commensal colonization. We detected both *B. fragilis* and *B. thetaiotaomicron* in the colons of the newborns 7 days after birth, demonstrating that both strains are transferred from the dam to her pups (Figure 4A and 4B). However, at 9 days of age, *B. fragilis* WT was present at a much higher proportion than *B. thetaiotaomicon* (Figure 4A), whereas Δ*nanH* was unable to compete with *B. thetaiotaomicron*, displaying a fitness defect (Figure 4B). Moreover, *B. fragilis* WT continued to dominate over *B. thetaiotaomicron* in 14 days old pups (Figure 4A), while Δ*nanH* was still unable to bloom at comparable levels (Figure 4B). Further, the defect in competitive fitness was retained by the Δ*nanH* during co-colonization with *B. thetaiotaomicron* of adult gnotobiotic mice as compared to *B. fragilis* WT (∼100-fold, *P*<0.05) (Figure 4C, 4D and Figure S6). These findings further emphasize the significance of NanH sialidase in conferring competitive fitness to *B. fragilis* during co-colonization of both newborn and adult mice with other *Bacteroides species*.

**Figure 4.**
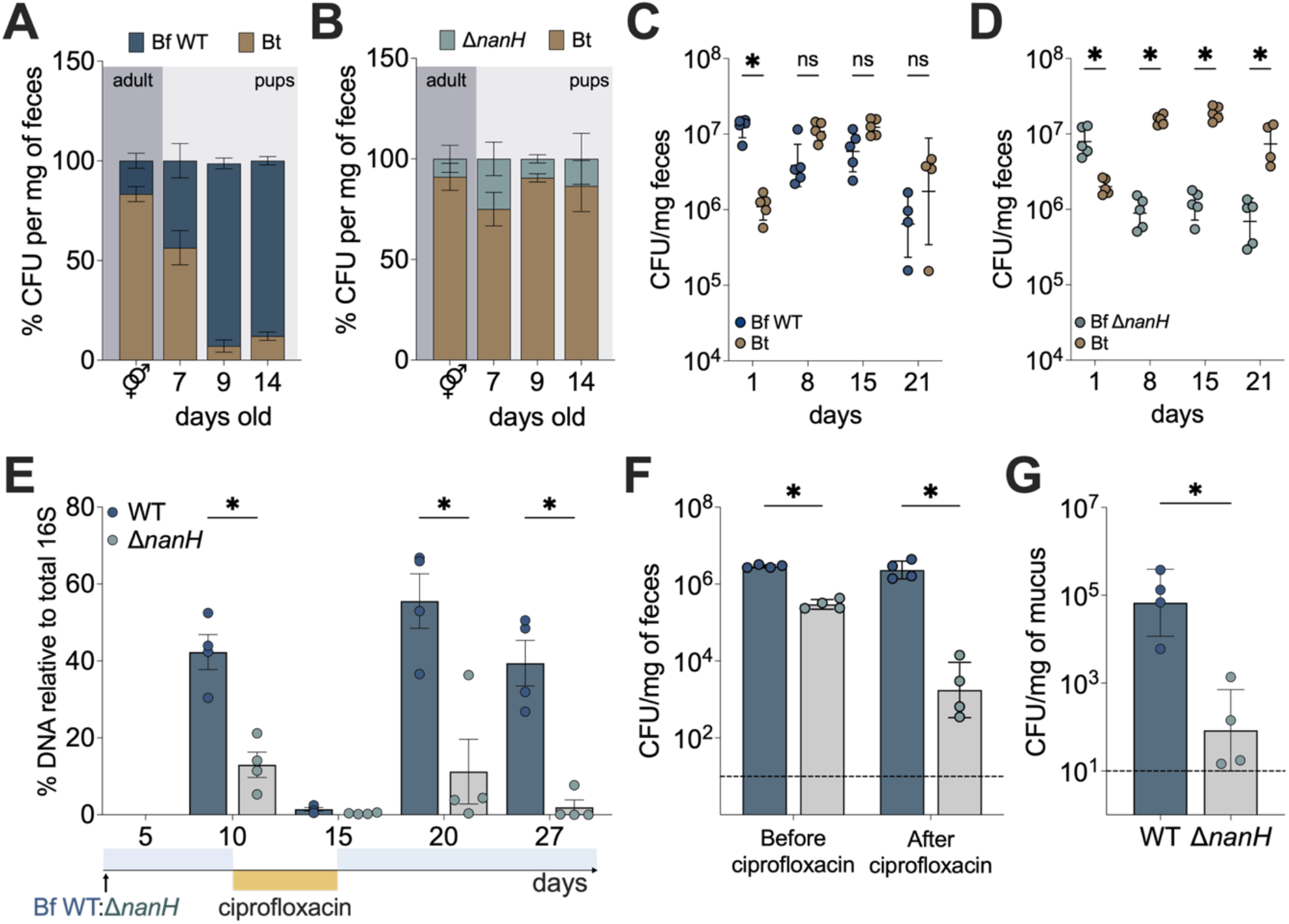
*B. fragilis* NanH facilitates commensal resilience and competitive fitness. (A) Abundance of *B. fragilis* WT and *B. thetaiotaomicron* in the feces of adult breeders (dark grey shaded area) and the feces of the newborn suckling pups (light grey shaded area) *n*≧3 per time point. (B) Abundance of Δ*nanH* and *B. thetaiotaomicron* in the feces of adult breeders (dark grey shaded area) and the feces of the newborn suckling pups (light grey shaded area) *n*≧3 per time point. (C – D) Germ free mice were inoculated with a single dose of either (C) *B. fragilis* WT (WT Bf) and *B. thetaiotaomicron* WT (Bt) or (D) Δ*nanH* and *B. thetaiotaomicron* WT at 1:1 ratio. Fecal CFUs were monitored weekly and determined by colony counts by plating on selective media and incubating anaerobically for 2 days. (E – G) Mice harboring the Simplified Human Intestinal Microbiota (SIHUMIx) were challenged with WT and Δ*nanH B. fragilis* (1:1) on day 0. The abundance of WT and Δ*nanH B. fragilis* was determined by qPCR relative to total fecal microbial 16S before (day 10), straight after (day 15), and during the recovery period (days 20 and 27) from the ciprofloxacin treatment. (F) CFU/mg of WT and Δ*nanH B. fragilis* in the feces before and after the ciprofloxacin treatment. (G) CFU/mg of WT and Δ*nanH B. fragilis* recovered from the colonic mucosa 13 days post ciprofloxacin treatment. *n* = 4. Fecal, colonic, and mucosal CFUs/mg were determined by colony counts and plating on selective media, using antibiotic resistance conferred by pFD430 plasmids. The dashed line defines the LOD = 10 CFU/mg. Data show geometric mean ± SD. \**P* < 0.05; \*\**P* < 0.01; \*\*\**P* < 0.001; \*\*\*\**P* < 0.0001; ns – not significant (Mann-Whitney test). Data represents 2 independent experiments.

In the past few decades, the widespread use of antibiotics has led to a substantial number of children receiving least one treatment of antibiotics before age two^38,39^. Consequently, antibiotic exposure disrupts the stability and composition of the microbiota, rendering individuals more susceptibility to various diseases^38^. We thus explored whether NanH mediates commensal resilience, enabling niche occupancy and recovery post antibiotic exposure in the context of a complex, defined microbial community. To test this, we orally gavaged adult mice harboring the Simplified Human Intestinal Microbiota (SIHUMIx)^40^ with *B. fragilis* WT and Δ*nanH* at 1:1 ratio. The mice were colonized for a 10-day period to allow for the stabilization of the *B. fragilis* strains into the established microbial community. We then introduced a broad-spectrum antibiotic, ciprofloxacin, in drinking water for 5 days, after which the mice were maintained on regular drinking water and the relative abundance of WT and Δ*nanH* was determined by qPCR and fecal colony counts to evaluate the ability of both strains to re-establish colonization after antibiotic perturbation. Consistent with our findings in co-colonized mice (Figure 2J), Δ*nanH* was present at a lower proportion than *B. fragilis* WT in mice harboring a complex bacterial community prior to the ciprofloxacin treatment (Figure 4E and 4F). Notably, while *B. fragilis* WT was able to recover from the ciprofloxacin treatment, Δ*nanH* was unable to re-establish colonization and gradually reduced both in the lumen (Figure 4E and 4F) and the intestinal mucosa (Figure 4G). These results demonstrate that the presence of NanH sialidase in *B. fragilis* is critical for the ability to re-establish its intestinal niche and compete in the simplified microbial community after antibiotic perturbation.

Several human cohort studies have provided evidence that a complex community of microbes is seeded from the mother during birth^12^. *B. fragilis* is a keystone species that can colonize the infant during birth and persist as a pioneering gut *Bacteroides* throughout an individual’s lifetime^12^. Interestingly, a metagenomic study characterizing the genetic repertoire of gut microbiomes in human populations revealed an enrichment of bacterial sialidases in infants younger than 6 months^41^. These observations suggest commensal sialidases may have significant impacts on the development of the gut microbiota during infancy and early life. To gain further insights into the prevalence and conservation of NanH sialidase among *B. fragilis* strains, we conducted a comprehensive analysis of over 300 publicly available *B. fragilis* genomes, including strains from pediatric donors. This revealed that NanH is a part of the *B. fragilis* core genome, with a high sequence identity (98%) among homologues across all *B. fragilis* strains (Table 2). These findings provide compelling evidence that the presence and functionality of NanH sialidase are conserved among *B. fragilis* strains, underscoring its importance in the colonization dynamics of this pioneer *Bacteroides* species. Indeed, supporting stable *B. fragilis* colonization during early life may play a vital role in maintaining immune tolerance in the gut^42,43^. We reveal that proportion of IL-10 producing FOXP3+ T_regs_ was significantly reduced in the mesenteric lymph nodes (MLNs) of mice mono-colonized with Δ*nanH* (Figure S7A), while the overall proportions of FOXP3+ T_reg_ remained comparable to *B. fragilis* WT (Figure S7B). Combined with our data indicating NanH mediates early life colonization, this might suggest that the stable niche occupancy is crucial for early life immune priming and the development of mature immune responses. Here, we demonstrate that sialylated HMOs received pre-weaning direct *B. fragilis* towards its intestinal niche, ensuring a stable and resilient colonization. This highlights the intricate interplay between HMOs, host glycans, and keystone bacterial species in shaping microbial composition during early life.

## DISCUSSION

Here, we report that HMOs from human milk play a key role in establishing intestinal colonization of commensal bacteria. Several studies have indicated a positive association between breastfeeding and the successful establishment of *Bacteroides* species in the infant gut^6,44^. Our findings unveil the molecular mechanisms directly linking HMO metabolism to stable colonization of a human commensal, *B. fragilis*. We demonstrate that in response to HMOs *B. fragilis* expresses enzymatic systems which facilitate their breakdown and concurrently function as mediators of intestinal colonization (e.g., NanH, CcfC, CcfD). Our data shows that defects in *B. fragilis* sialidase, NanH, result in its impaired growth on HMOs and reduced intestinal colonization in early life, indicating that these processes are intricately interlinked.

Previous studies have reported that the gut microbiome of vaginally delivered infants were dominated by *Bacteroides* species^45–47^. In contrast, infants born via C-section tend to exhibit a low *Bacteroides* signature^3,4^. Further analyses revealed that while the microbiota of both vaginal and C-section delivered infants were similar in the first week of life, C-section infants lost *Bacteroides* species after week two of life^3^. It has been reported that mothers who undergo C-sections may have delayed initiation of breastfeeding^48,49^. Based on our findings – that HMOs induce the colonization program of *B. fragilis* – we speculate whether *Bacteroides* species are maintained in breastfed infants due to induction of bacterial genes associated with stable intestinal colonization, such as the *B. fragilis* sialidase, NanH. Furthermore, an analysis of HMOs from postpartum mothers in Malawi revealed that mothers whose infants were severely stunted had significantly lower levels of sialylated HMOs^50^. Transplantation of the fecal microbiota of malnourished Malawian infants into germ-free mice maintained on a diet of sialylated oligosaccharides from bovine milk improved growth and development in a microbiome-dependent manner. Indeed, out of all microbial species within the community, *B. fragilis* exhibited the most prominent transcriptional response to sialylated oligosaccharides from bovine milk. The upregulated *B. fragilis* genes included the Nan operon and a pair of SusC/D proteins (BF1804/BF1805) from HMO PUL 1, adjacent to NanH (BF1806) we describe in our study (Figure 1). In contrast, no significant changes were detected in any of the *Bifidobacterium* (*B. catenulatum*, *B. breve* and *B. bifidum*) or other *Bacteroides* strains. Further, NanH was previously shown to release sialic acid from submaxillary mucins^51^, supporting its potential role in the deconstruction of colonic mucins. Interestingly, *B. thetaiotaomicron* possesses a PUL homologous to the HMO PUL 1, in which proteins have been shown to display activity against GlcNAc-β1-3-Gal and β1-4-Gal decorations present in complex N-glycans^36^ and also comprising type I and II HMOs. This suggests that, in addition to HMOs and mucus glycans, enzymes in *B. fragilis* HMO PUL 1 might provide access to a variety of host glycoproteins. Altogether, we propose a model where sialylated HMOs act as the signal for a pioneer gut commensal to occupy the mucosal niche at early life, after which structurally similar mucin glycans maintain a bacterial reservoir, supporting commensal persistence and resilience after weaning.

Furthermore, we demonstrate that the genes induced by HMOs confer *B. fragilis* a competitive advantage during early life and are essential for stable colonization in the context of simple (mono- and co-colonization) and complex microbial communities. Our study here reveals the involvement of NanH-mediated colonization in the recovery and resilience of *B. fragilis* following antibiotic treatment. Importantly, a significant portion of children receive at least one course of antibiotics during the first two years of life^38^. Given the ubiquitous presence and conservation of NanH in all *B. fragilis* strains, including those from pediatric populations (Table 2), this supports its functional importance in the co-evolutionary relationship between *B. fragilis* and its mammalian host. Recent work has shown a correlation between reduced levels of a sialylated milk oligosaccharide (DSLNT) and the development of inflammatory disorders such as necrotizing enterocolitis (NEC) in breastfed infants^52,53^. Our data indicates that mono-colonization with *B. fragilis*, lacking NanH, leads to reduced proportions of mesenteric FOXP3+ IL-10+ T_regs_, further pointing to the significance of sialylated HMOs in maintaining *B. fragilis* as a stable pioneer species during early life and emphasizing its crucial role in promoting host immunity and health^43,54,55^. Future studies will examine the immunological consequences of *B. fragilis* colonization during this critical developmental window in the first weeks of life. We speculate that other commensals likely employ analogous strategies, whereby HMO metabolism triggers the expression of colonization mediators, perhaps extending beyond the classes of microbial sialidases described in this study. Collectively, our findings demonstrate that the mammalian host curates its indigenous microbiota by providing specific substrates, such as HMOs, within a developmental window to ensure stable colonization of beneficial microbes.

## AKNOWLEDGEMENTS

We thank members of the Chu lab for technical support and helpful discussions. We also thank G. Donaldson and A. Khosravi for valuable feedback and discussions. This work was supported by grants from the National Institute of Diabetes, Digestive and Kidney Diseases (NIDDK) R00 DK110534 and P30 DK120515, and a Seed grant made available through the UC San Diego Larsson-Rosenquist Foundation Mother-Milk-Infant Center of Research Excellence to H.C. L.B. is UC San Diego Chair of Collaborative Human Milk Research endowed by the Family Larsson-Rosenquist Foundation (FLRF), Switzerland. Additional support was provided to H.C. by the Chiba University-UC San Diego Center for Mucosal Immunology, Allergy and Vaccines (cMAV), CIFAR Humans and the Microbiome Program, and The Hartwell Foundation. Support to L.R. was provided by the UCSD Graduate Training Program in Cellular and Molecular Pharmacology through an institutional training grant from the National Institute of General Medical Sciences, T32 GM007752.

## METHODS

### Bacterial strains and growth conditions

Bacterial strains used in this study are listed in Table 3. Type strains (ATCC) were used for growth experiments unless otherwise stated. *Bacteroides* strains were grown in BHI-S (Brain Heart Infusion, BD) medium supplemented with 0.5 % hemin and 0.5 µg/ml vitamin K (Sigma Aldrich) for 16 h at 37 °C under anaerobic conditions (10 % H_2_, 10 % CO_2_, 80 % N_2_; Coy Lab Products). For growths utilizing a sole carbon source, ZMB1 (Zhang-Millis-Block1)^56^ defined medium was supplemented with 0.5 % hemin and 0.005 % yeast extract (Sigma Aldrich). Overnight cultures were sub-cultured into fresh BHI-S and grown to OD_600_=0.6-0.8, pelleted by centrifugation at 11,000 g for 1 min, washed with sterile ZMB1 twice and resuspended in ZMB1. Cultures were then normalized to the same OD_600_ inoculated at 2.5% into ZMB1, containing 15 mg/ml pooled mixed, pooled acidic, pooled neutral HMOs or 5 mg/ml single HMOs (2’FL, 3FL, 6’SL, LNnT, or LNT) as a sole carbon source. Growth assays were performed in triplicates in a 96-well plate using Biotek Synergy HT plate reader. All growth curves are representative of at least three independent experiments. Where appropriate gentamicin (200 µg/ml), erythromycin (10 µg/ml), ampicillin (100 µg/ml), chloramphenicol (10 µg/ml), or tetracycline (6 µg/ml) were used for bacterial selection.

### Generation of deletion mutants and pFD340-conjugates

Deletions of BF1806 (NanH) was generated by amplifying 1 kb fragments upstream and downstream of the region (Table 4) and cloned into pKNOCK-bla-erm^57^ using NEBuilder (NEB). The plasmid was conjugally transferred into *B. fragilis* NCTC9343 or *B. fragilis* ΔCCF using *E. coli* S-17 *λ-pir*. Conjugates were selected based on erythromycin resistance and the second recombination event was encouraged by daily passage until erythromycin resistance was lost. Scarless inframe deletions were confirmed by PCR using primers listed in Table 4 and sequencing (Primordium).

To create antibiotic resistant strains used for colonization experiments, *E. coli* S-17 *λ-pir* were first transformed with pFD340-*tetQ* (tetracycline resistant, TetR) and pFD340-*cat* (chloramphenicol resistant CmR) plasmids^17^. These were then conjugally transferred to generate *B. fragilis* WT+pFD340-*tetQ*, *B. fragilis* Δ*nanH*+pFD340-*cat*, *B. thetaiotaomicron*+pFD340-*tetQ*, *B. thetaiotaomicron*+pFD340-*cat*. The conjugates were selected based on erythromycin (10 mg/ml) resistance and secondary resistance to tetracycline (6 mg/ml) or chloramphenicol (10 mg/ml). During in vitro culturing, erythromycin was added to rich media (BHI-S) to ensure plasmid maintenance.

### HMO isolation from pooled donor human milk

Pooled HMOs (pHMOs) were isolated from pooled donor human milk as previously described^58^. After centrifugation, the lipid layer was removed, and proteins were precipitated from the aqueous phase by addition of ice-cold ethanol and subsequent centrifugation. Ethanol was removed from the HMO-containing supernatant by roto-evaporation. Lactose and salts were removed by gel filtration chromatography over a BioRad P2 column (100 cm x 316 mm, Bio-Rad) using a semi-automated fast protein liquid chromatography (FPLC) system. pHMO composition was measured as described below. Only pHMOs with less than 2 % lactose were used for bacterial utilization experiments.

### HMO analysis

Analysis of HMOs in defined media was performed by high-performance liquid chromatography (HPLC) with fluorescence detection as previously described for human milk^59^ with slight modifications. 10 µL of media was spiked with 12 ng/ml of maltose as an internal standard, lyophilized, and directly labeled with the fluorophore 2-aminobenzamide (2AB). 2AB-labeled HMOs were separated on a TSKgel Amide-80 column (2.0 mm ID x 15 cm, 3m, Tosoh Bioscience) and detected at 360 nm excitation and 425 nm emission. HMO peaks were annotated based on standard retention times and quantified in reference to the internal standard. HMO utilization at time x was calculated in reference to HMO concentrations in the media at the beginning of each experiment (t=0).

### Thin layer chromatography

Aliquots of 3-12 μl of cell-free supernatant or enzymatic reactions were spotted onto silica coated plates (Merck, TLC Silica gel 60 F254) and resolved in butanol:acetic acid:water (2:1:1). Sugars were visualized with diphenylamine (DPA) stain^60^. 1 mM of each glucose, galactose, fucose, and lactose were run alongside samples and used for reference.

### RNA extraction, cDNA synthesis and relative gene expression

Overnight culture of *B. fragilis* was inoculated into defined medium containing 15 mg/ml pHMOs and 10 mg/ml glucose (Sigma Aldrich). Bacterial pellets were collected from mid-exponential phase (OD_600_=0.6-1.0), resuspended in RNA later (Thermo Scientific), and stored at -80 °C. For the determination of the transcript levels during mono-colonization, colonic contents or the mucosal scrapings were placed in RNA later immediately after collection and stored at -80 °C. RNA was extracted from bacterial pellets, fecal material, or the mucosal lining using a NucleoSpin RNA isolation kit (Macherey-Nagel), 0.5-1 μg was reverse transcribed into cDNA (SuperScript IV VILO Master Mix, Thermo Scientific) as per manufacturer’s instructions, and diluted 1:5. Transcription levels were assessed by qPCR (SYBR green qPCR Master Mix, Life Technologies) in Quantistudio 5 (Life Technologies) and normalized to gyrase B using primers listed in Table 4. For the in vitro experiments relative expression was compared to glucose grown cells.

### Quantitative Multiplex and Label-Free Proteomics

Peptide preparation: Overnight cultures of *B. fragilis* NCTC 9343 was inoculated into ZMB1 media containing 15 mg/ml pHMOs or 10 mg/ml glucose as a sole carbon source. All cultures were performed in triplicates. 4 ml of culture was collected at early exponential (OD_600_ = 0.5), mid-exponential (OD_600_ = 1.0), and late exponential (OD_600_ = 1.5) growth phases, pellet was washed in sterile PBS three times, and stored at -20 °C prior to analysis. Pellets were resuspended in lysis buffer (6M urea, 7% SDS, 50 mM TEAB, titrated to pH 8.1 with phosphoric acid) with protease and phosphatase inhibitors added (Roche, CO-RO and PHOSS-RO) and sonicated. Proteins were reduced, alkylated, then trapped using ProtiFi S-Trap columns (ProtiFi, C02-mini), digested with sequencing-grade trypsin (Promega, V5113), and eluted according to manufacturer protocols. Eluents were desalted on SepPak C18 columns (Waters, WAT054960). Peptides were quantified using a Pierce Colorimetric Peptide Quantification Assay Kit (Thermo Scientific, 23275). 50 μg of each sample was separated for multiplex proteomic analysis, with several samples aliquoted twice to serve as technical duplicates and fill multiplex channels.

### Labeling and fractionation

The labeling scheme for multiplex experiment is included in the Supplemental Material. Samples were labeled using Tandem Mass Tag (TMT) 16plex reagents (Thermo Scientific, A44520; lot number XA341491) following manufacturer protocol, then combined into a single multiplex. The plex was desalted on SepPak C18 columns and dried under a vacuum. The plex was then fractionated using reverse phase high pH liquid chromatography on a 10-40% acetonitrile (ACN) gradient to increase sequencing depth, as previously described. The resulting 96 fractions were concatenated into 24 fractions by combining alternating wells within each column, and 12 alternating fractions were used for mass spectrometry analysis^61^

### LC-MS/MS

One μg of each fraction was loaded and analyzed on an Orbitrap Fusion Tribrid mass spectrometer with an in-line Easy-nLC 1000 System and an in-house pulled and packed column, as previously described^61^. Peptides were eluted after loading using a gradient ranging from 6% to 25% ACN with 0.125% formic acid over 165 minutes at a flow rate of 300 nL/min. Data were acquired in data-dependent mode with polarity set to positive. MS1 spectra were acquired in the Orbitrap with a scan range of 500-1200 m/z and a mass resolution of 120,000. Ions selected for MS2 analysis were isolated in the quadrupole and detected in the ion trap. MS2 ions were fragmented with high-energy collision– induced dissociation, and MS3 fragment ions were analyzed in the Orbitrap. All data acquired were centroided.

### Data processing and normalization

Raw files were processed using Proteome Discoverer 2.5. Using the SEQUEST algorithm, MS2 data from *B. fragilis* NCTC 9343 was queried against Uniprot proteome UP000006731, downloaded May 2022. Resulting peptide spectral matches were filtered at a 0.01 FDR by the Percolator module against a decoy database. Peptide spectral matches were quantified using MS3 ion intensities, exported, then summed to the protein level. Entire plex was then batch corrected in a multistep process as previously described^62^. Lastly, the data were normalized by log_2_ transformation and technical duplicates were averaged. Final data represent normalized relative protein abundances.

### Protein differential abundance

The normalized relative protein abundance matrices were imputed in R for differentially abundance testing. A Welch’s t-test was conducted for each protein between the given time point and the glucose set to find proteins with change in differential abundance in relation to the control. Benjamini-Hochberg correction for multiple testing was applied to the resulting p-values. Proteins were considered significant if the corrected p-value was less or equal to 0.05 and the log-fold change was greater than or equal to 2. This data was used to create volcano plots in Prism. An ANOVA test between all timepoints was conducted, and significant protein hits relative to glucose (LFC ≧2, Bonjamini-Hochberg corrected p-value ≦ 0.05) were plotted in a heatmap in R.

### Gnotobiotic mouse experiments

4-week-old sex matched germ-free mice were C57BL/6J mice (JAX: 000664) were orally gavaged with wild type (WT) and Δ*nanH B.fragilis* (8E07 CFUs per dose). For experimental purposes, upon colonization mice were housed in sterile, sealed positive pressure cages with double HEPA filtration (Allentown) with autoclaved chow (Lab Diet 5010) and drinking water. For the co-colonization experiments, mice were orally gavaged with bacterial suspension containing equal proportions of WT*B. fragilis* and the isogenic Δ*nanH* mutant, WT*B. fragilis* and WT *B. thetaiotaomicron*, or Δ*nanH B. fragilis* and WT *B. thetaiotaomicron* (∼8x10E7 CFUs of each strain per dose). For the sequential colonization experiments, 4–5-week-old germ-free C57BL/6J mice were mono-colonized with the initial strain (∼8x10E7 CFUs per dose) for 7 days, and the challenge strain was introduced by oral gavage on day 8. All strains used for colonization experiments retained pFD340 plasmids, conferring resistance to erythromycin and either tetracycline or chloramphenicol. Gentamicin (100 μg/ml) and erythromycin (10 μg/ml) were added to the drinking water to ensure plasmid maintenance. At each time point, a fresh fecal pellet was collected, resuspended in sterile PBS, serially diluted, and plated on BHI-S agar containing either tetracycline (6 μg/ml) or chloramphenicol (10 μg/ml) to allow for differentiation between *Bacteroides species*. Abundance of each strain was enumerated by CFU per milligram of feces. All procedures were performed in accordance with the guidelines and approved protocols from the IACUC of UC San Diego.

### Experiments in mice colonized with a defined microbial community

The C57BL/6J mice were colonized with the Simplified Human Intestinal Microbiota (SIHUMIx) composed of 7 bacterial species: *Anaerostipes caccae* (DSMZ 14662), *Bifidobacterium longum* (NCC2705), *Blautia producta* (DSMZ 2950), *Clostridium bytiricum* (DSMZ 10702), *Clostridium ramosum* (DSMZ 1402), *Escherichia coli* K-12 (MG1665), *Lactobacillus plantarum* (DSMZ 20174) and bred in our lab. Age and sex matched 5-6 weeks old mice were then inoculated with a single gavage of bacteria mixture of WT and Δ*nanH B. fragilis* (8E07 CFUs of each strain) and the strains were allowed 10 days for the integration into the community. On day 10, mice were then treated with 0.625 mg/ml of ciprofloxacin (Sigma Aldrich) in drinking water for 5 days, after which they were switched back and maintained on regular water for further 13 days. At each time point, bacterial CFUs were determined by plating on selective media and pFD340-based differentiations. In addition, total microbial genomic DNA was extracted using Quick DNA Fecal/soil Microbe miniprep kit (Zymo Research) as per manufacturer’s protocol. The relative abundance of WT and Δ*nanH B. fragilis* was assessed by absolute qPCR using strain specific and universal 16S primers listed in Table 4.

### Mucosal scraping for bacterial quantification

Colons were excised and washed with sterile ice-cold PBS to remove luminal contents. The colons were then cut open longitudinally, washed until fecal material was no longer visible, and the mucus was scraped with light pressure using the blunt side of the sterile tweezers. Collected mucus was placed in pre-weighed tubes, containing 500 μl of sterile PBS, and homogenized using sterile stainless-steel beads (⌀3.2mm) and a bullet blender (Next Advance) for 3 mins at speed 8. Homogenized mucus was then serially diluted and plated on BHI-S with appropriate antibiotic for enumeration by colony counts.

### Vertical transfer experiments and colonization of suckling pups

Breeding pairs of 8-week-old male and female germ-free *Rag1*-/- (JAX:008449) mice were orally gavaged with a bacterial culture comprising of WT and Δ*nanH B. fragilis*, WT *B. fragilis* and WT *B. thetaiotaomicron*, or Δ*nanH B. fragilis and* WT *B. thetaiotaomicron* (8E07 CFUs of each strain per dose). The breeders were housed in sterile, sealed positive pressure cages with double HEPA filtration (Allentown), with autoclaved chow (Lab Diet 5010) and drinking water containing gentamycin (100 mg/ml) and erythromycin (10 mg/ml). Upon delivery of the litter, the pups were sacrificed at 7, 9-10, and 14 days of age, colons were excised, and the contents were resuspended in 300 μl of sterile PBS in pre-weighed tubes. The contents were then homogenized with sterile stainless-steel beads (⌀3.2mm) in a bullet blender (Next Advance) for 3 mins at speed 8. The homogenized colonic material was then serially diluted and plated on either tetracycline or chloramphenicol containing BHI-S plates to allow for differentiation between the competing species. CFUs/mg for each pup were calculated from colony counts and presented as a ratio relative to the breeders.

### Stomach processing for analysis by TLC or HPLC

The stomachs were collected from pups of 0-1, 2, 3, and 4 weeks of age and placed in 1 ml of sterile PBS. The tissue was homogenized using stainless-steel beads (⌀3.2mm) and a bullet blender (Next Advance) for 5 mins at speed 10. Homogenized tissue was then centrifuged at 14,000 x g for 3 mins, the supernatant containing soluble glycans was separated and used for the downstream analysis.

### Isolation of cells from tissues

Mesenteric lymph nodes (MLN) were processed by mashing tissues through 100 µm cell strainer (BD Falcon) to generate single cell suspensions.

### Flow cytometry

Cells were stained for 30 min at 4 °C with either LIVE/DEAD fixable violet or yellow dead stain Kit (Life Technologies), with empirically titrated concentrations of the following antibodies: eF450-conjugated anti-mouse CD4 (clone: RM4-5), and LIVE/Dead Fixable Yellow Dead Cell Stain Kit (Life Technologies). For intracellular staining, cells were fixed and permeabilized using the Foxp3/Transcription Factor Staining Buffer Set (eBioscience) according to the manufacturer’s protocol. Intracellular staining was performed with the following antibodies: PE-conjugated anti-mouse IL-10 (clone: JES5-16E3), and APC-conjugated anti-mouse Foxp3 (clone: FJK-16s) for 3-4 hours. All antibodies were purchased from Thermo Scientific/eBiosciences, BD, and Biolegend. Cell acquisition was performed on LSRFortessa (BD), and data was analyzed using FlowJo software suite (TreeStar).

### Quantification of *Bacteroides* by absolute qPCR

The presence of each *Bacteroides* species was determined by absolute qPCR (SYBR green qPCR Master Mix, Life Technologies) from the total genomic DNA. Species specific primers listed in Table 4 were used to determine abundance of WT and Δ*nanH B. fragilis* and *B. thetaiotaomicron*. Each gDNA sample was diluted to 20 ng/µl and the relative abundance of the species was determined from standard curves prepared from gDNA extracted from pure overnight cultures (range 20 ng, 10 ng, 1 ng, 0.1, ng, 0.01 ng, 0.001 ng) or a fecal pellet collected from SIHUMIx colonized mice without *B. fragilis* for the bacterial 16S (80 ng, 45 ng, 20 ng, 10 ng, 1 ng, 0.1 ng, 0.001 ng). Data was analyzed in Quantstudio 5 (Life Technologies).

### Data availability

Proteomic data and additional supplementary files for reanalyzing the data collected here are available online at https://massive.ucsd.edu under study ID MSV000090386. https://github.com/rolesucsd/hmo_proteomics

### Bioinformatics analysis

Glycoside hydrolases and PUL boundaries were identified using Cazy database and PUL-DB^28^. Modular organization of proteins was assessed in InterPro^63^. Sequence alignments and percent identities were determined in Clustal Omega^64^. The nature of the N-terminal signal peptides was identified in SignalP 5.0^65^. The phylogenetic tree for NanH was created by running BlastP against the non-redundant protein database and selecting the top hit for the species of interest. The alignment was created by running Clustal Omega with the *B. fragilis* NanH as well as the other protein hits. The phylogenetic tree was created by Neighbor-Joining and plotted in R.

