## Supplemental Figures for "A bacterial sialidase mediates early life colonization by a pioneering gut commensal"

### FIGURE S1

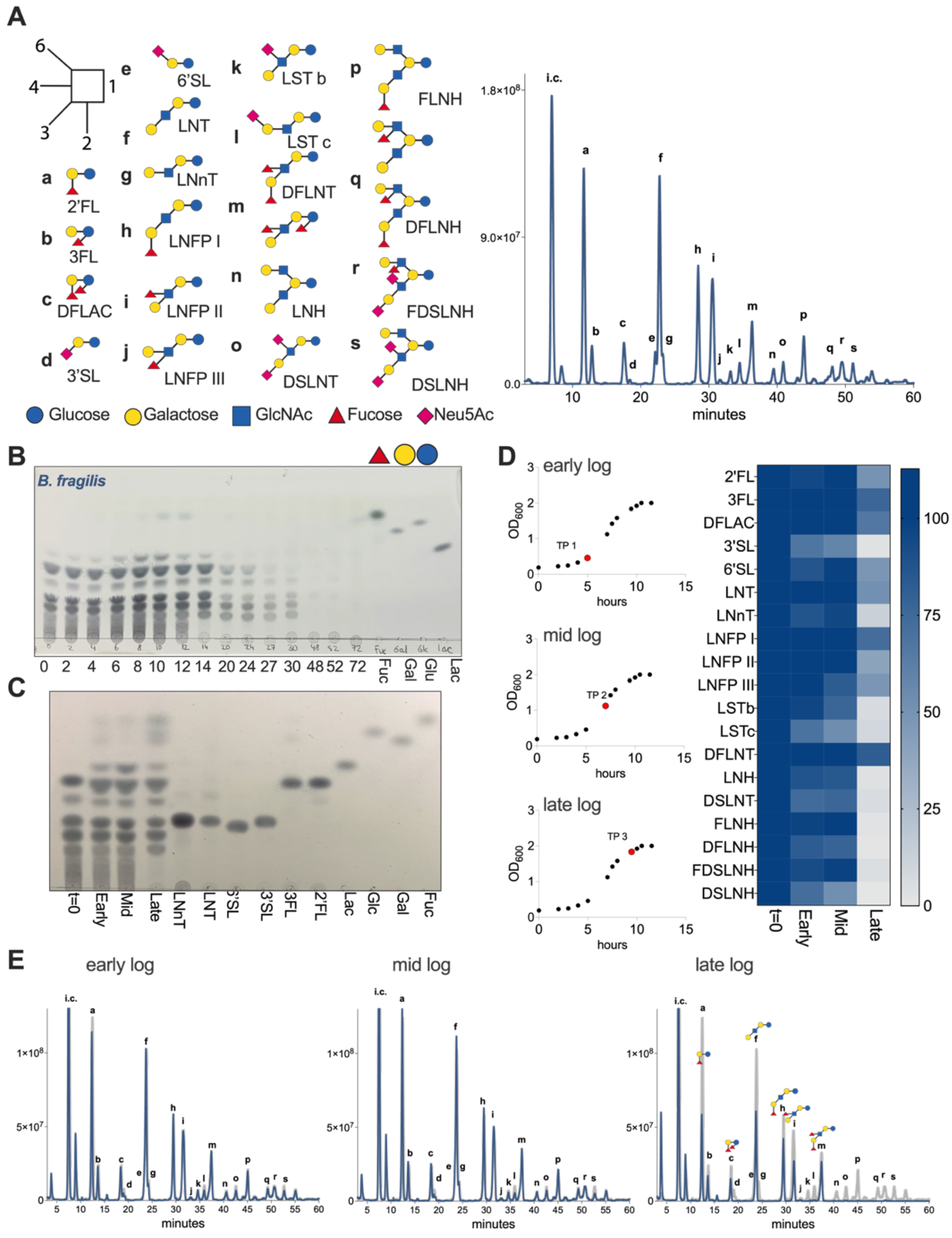


**Figure S1. Biphasic utilization of HMOs by *B. fragilis***

(A) Composition of the pooled HMOs present in the donor milk samples used in the study. Diagrams of 19 most abundant HMOs are shown on the left, HPLC chromatogram of the sample is shown on the right. Alphabetic labelling of individual HMOs corresponds to the peaks shown in the HPLC chromatogram. Maltose (2 ng/ml) was used as internal control (i.c). Sugars are color coded as indicated, linkage schemtics is displayed in the top right corner.

(B) TLC analysis of the cell free supernatant taken over the course of *B. fragilis*' growth on pHMOs.

(C) TLC analysis of the supernatant taken from early-, mid-, and late- log phases of *B. fragilis* growth on pHMOs. Standards include (1 mM): Lacto-neo-N-tetraose (LNnT), lacto-N-tetraose (LNT), 6'-sialyllactose (6'SL), 3'-sialyllactose (3'SL), 3-fucosyllactose (3FL), 2'-fucosyllactose (2'FL), Lactose (Lac), Glucose (Glc), Galactose (Gal), Fucose (Fuc).

Growth of *B. fragilis* on 15 mg/ml pHMOs, whole cells were collected at indicated time points (TP) (red circle) for analysis by comparative proteomics.

(D) HPLC analysis (right) of the supernatant collected from indicated growth phases (left). Relative abundance of each glycan was determined relative to the internal control.

(E) HPLC chromatograms of the supernatant from indicated growth phases. Peaks are labelled alphabetically to correspond to structures shown in panel A. Maltose (2 ng/ml) was used an internal control (i.c).

**Figure S2.**

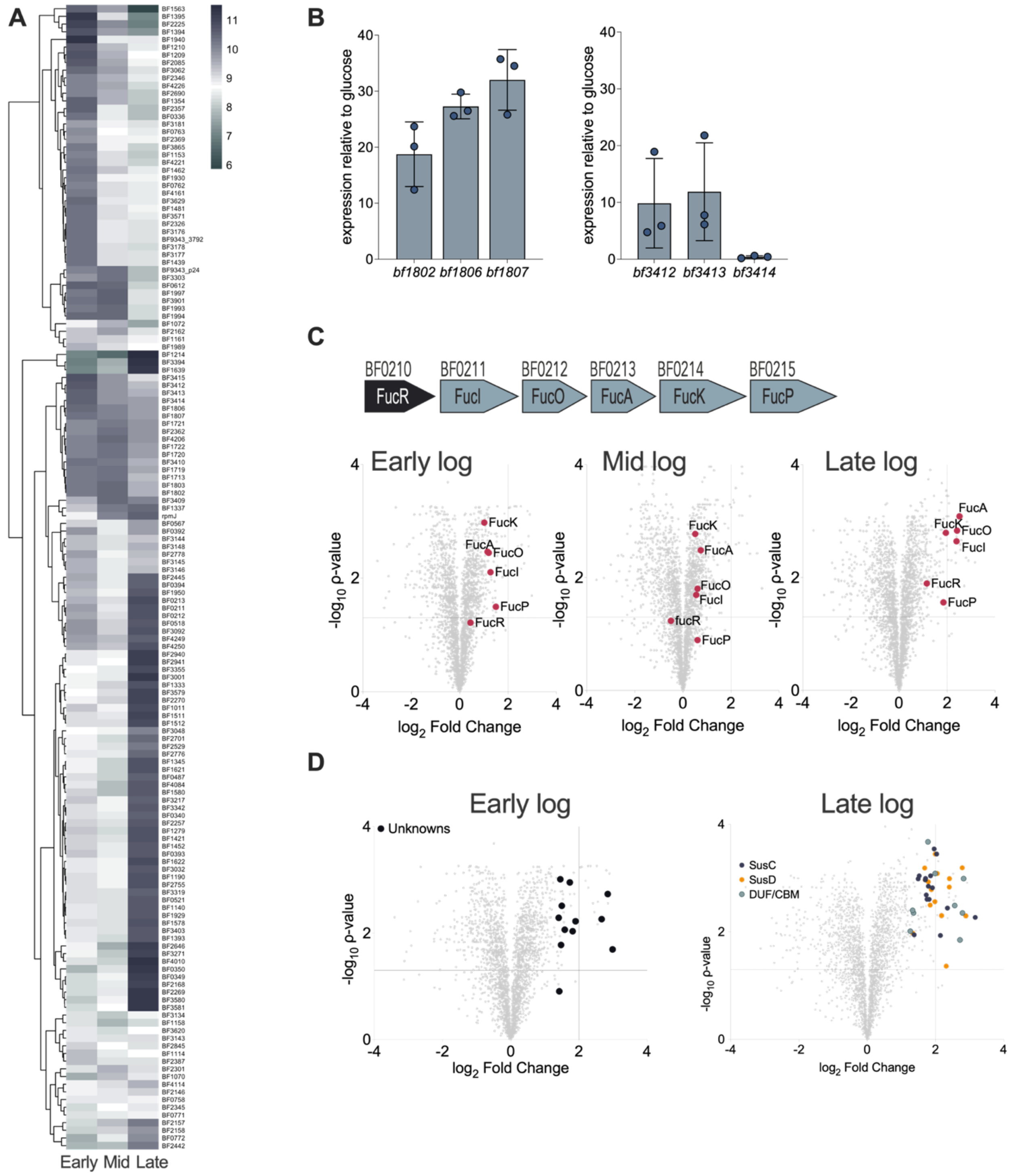


**Figure S2. Comparative proteomic analysis and relative expression of genes during *B. fragilis* growth on pHMOs**

(A) Abundance of proteins in HMO-grown *B. fragilis,* relative to glucose. Heatmap shows proteins with a max $\log_{10}$ fold change ≥2 and a Benjamini Hochenburg corrected *P*-value

< 0.05 between samples across three growth phases, early-, mid-, and late-log. Hierarchical clustering was performed using Euclidean distance. Scale bar shows highly abundant proteins in purple, lower abundant in green.

(B) Fold change relative to glucose of *bf1802*, *bf1806* (*nanH*), *bf1807* encoding for SusC, GH33, and GH20 in the HMO PUL 1, respectively, in comparison to *bf3412*, *bf3413*, and *bf3414* composing the Don operon. Transcript levels were assessed by qPCR from pHMO grown *B. fragilis* and normalized to glucose ($n$ = 3 cultures per substrate).

(C) Relative abundance of proteins composing the FucRIOAKP during growth on pHMOs. Organization of FucRIOAKP operon in *B. fragilis* is shown on top, FucR is a regulator, FucI - isomerase, FucO - reductase, FucA - aldolase, FucK - fuculokinase, FucP - permease.

(D) $Log_2$ fold change in pHMOs vs glucose *B. fragilis* during early- and late-log phase. Proteins of unknown function are shown in black circles, proteins in purple show SusCs, orange - SusDs, 13 pairs of SusC/Ds and 2 lone SusDs were significantly abundant during late-log growth. Proteins highlighted in green contain domains of unknown function (DUF) or carbohydrate binding modules (CBMs).

**Figure S3.**

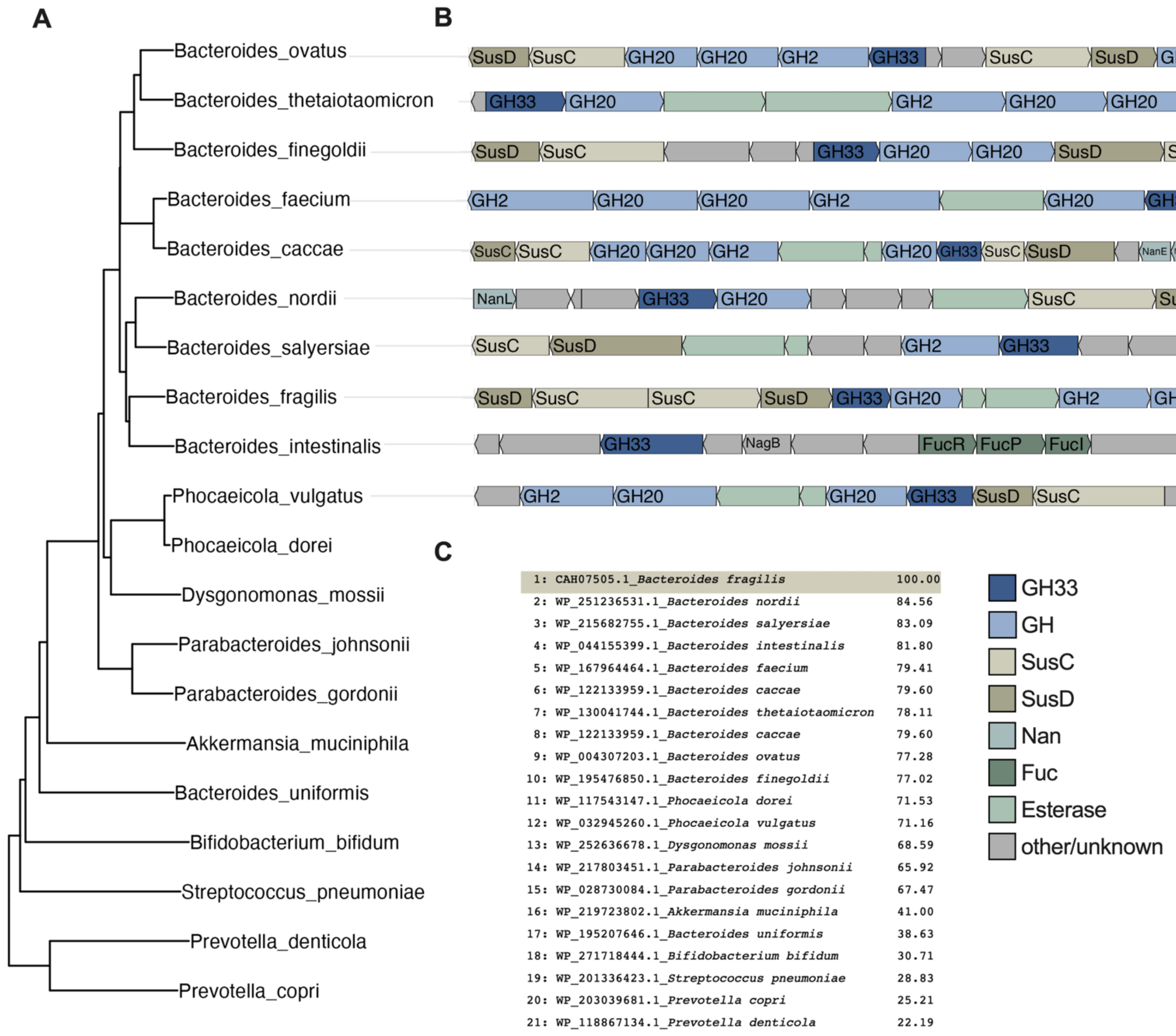


**Figure S3. Phylogenetic analysis of GH33 family members**

(A) Phylogenetic tree of GH33s present in different microbial species. The sequences were retrieved by BlastP and the tree was generated to scale by Neighbor-Joining in R.

(B) Organization of PULs enclosing GH33 homologs among *Bacteroides species*. Protein orientation was retrieved from the Integrated Microbial Genomes (IMG) database.

(C) Percent identity of proteins in the GH33 family relative to BF1806 (NanH), matrix was generated in Clustal Omega.

**Figure S4.**

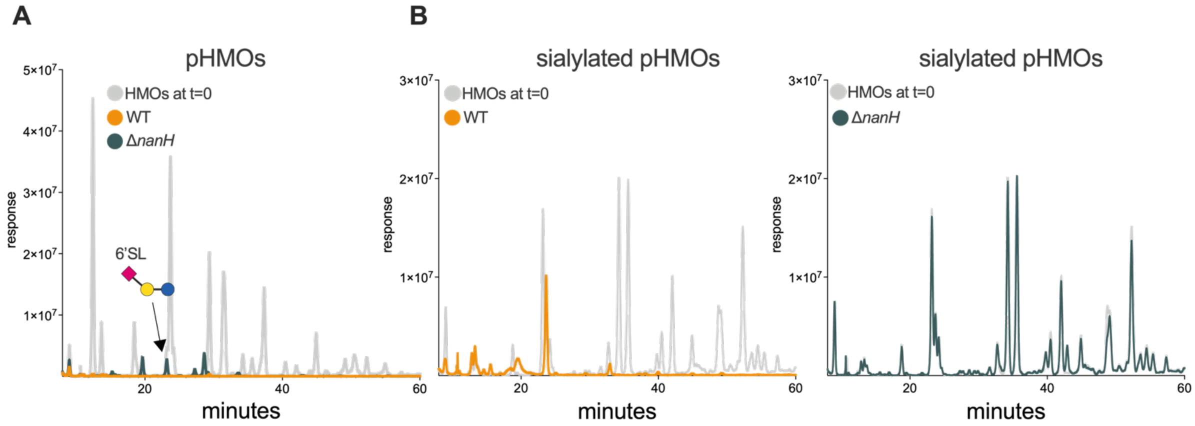


**Figure S4. Growth of WT and mutant *B. fragilis* on pooled and sialylated HMOs**

(A) Bacteria were grown anaerobically in defined media containing 15 mg/ml of substrate for 48 hours**.** HPLC analysis of the supernatant collected after growth of WT and Δ*nanH B. fragilis* on mixed pHMOs.

(B) HPLC analysis of the supernatant collected after growth of WT and Δ*nanH B. fragilis* on pooled sialylated HMOs. Grey trace shows substrate before bacterial growth, yellow trace - WT *B. fragilis*, green - Δ*nanH B. fragilis*.

**Figure S5.**

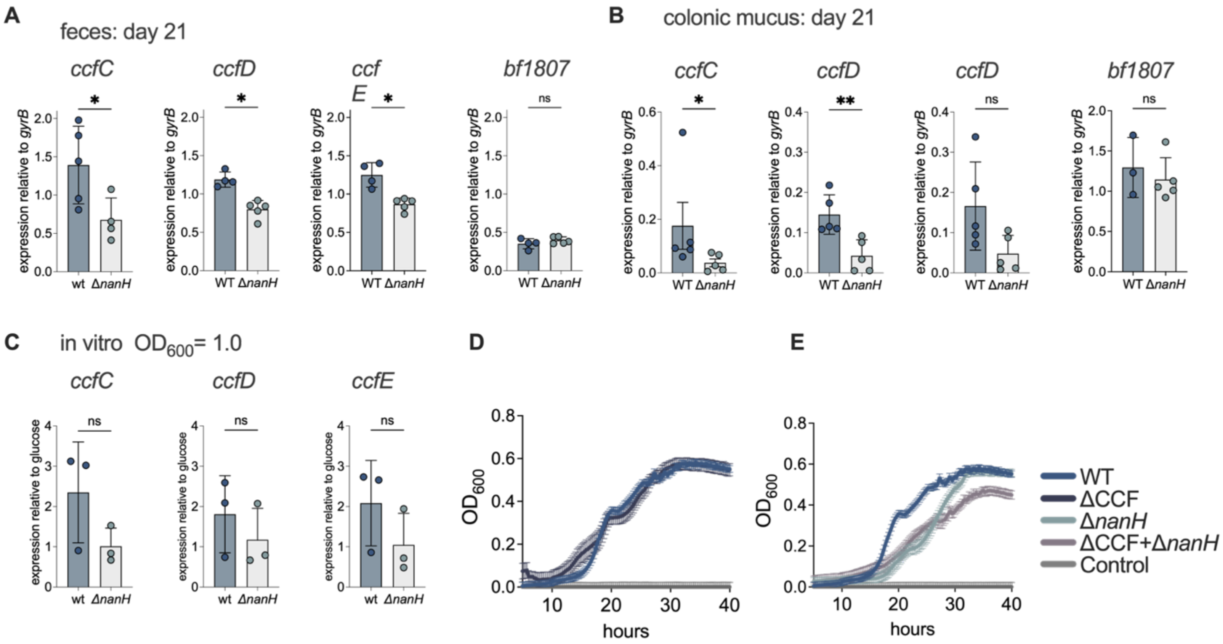


**Figure S5. Relative expression of the CCF locus during mono-colonization with WT and Δ*nanH B. fragilis***

(A – B) Relative expression of *ccfC* (*bf3581*), *ccfD* (*bf3580*), and *ccfE* (*bf3579*) in fecal content (A) and colonic mucosa (B) of mice mono-colonized with WT or Δ*nanH B. fragilis* for 21 days. RNA was extracted from the colon contents of mono-colonized mice, expression levels were assessed by qPCR and normalized to *gyrB* ($n$ = 4-5 mice per group), expression level of *bf1807* (GH20) was used as a control.

(C) Relative expression of *ccfC* (*bf3581*), *ccfD* (*bf3580*), and *ccfE* (*bf3579*) in WT and Δ*nanH B. fragilis* during growth on pHMOs *in vitro*. Bacterial cultures were grown to $OD_{600}$=1.0 in 15 mg/ml of pHMOs, transcript levels were normalized to *gyrB* and glucose-grown *B. fragilis* ($n$ = 3 cultures per group). Non-parametric Mann-Whitney test: *$P < 0.05$; **$P < 0.01$; ns – not significant.

(D – E) Growth of WT; ΔCCF, Δ*nanH*, and Δ*nanH*ΔCCF *B. fragilis* on 10 mg/ml pHMOs. Solid line shows the mean of three biological replicates ± SD.

**Figure S6.**

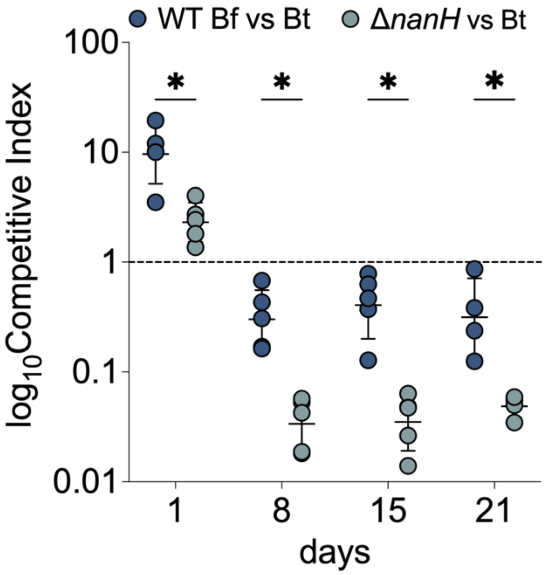


**Figure S6. NanH sialidase is required for competitive fitness in adult mice**

(A) The competitive index (C.I) between *B. fragilis* WT and *B. thetaiotaomicron* (blue circles); Δ*nanH* and *B. thetaiotaomicron* WT (green circles) during 1:1 co-colonization of gnotobiotic mice. The C.I demonstrates the ratio between the output and the initial inoculum.

**Figure S7.**

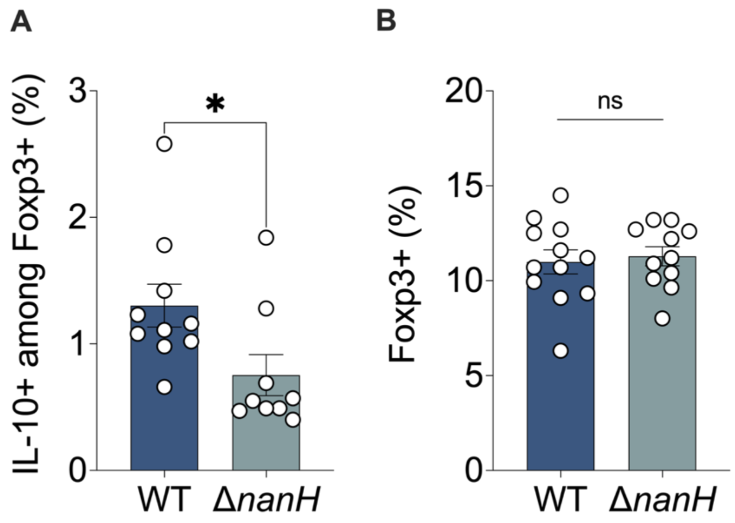


**Figure S7. Induction of CD4+Foxp3+IL-10+ Tregs in mesenteric lymph nodes during mono-colonization with WT and Δ*nanH B. fragilis***

(A) Proportion of IL-10 producing FOXP3+ cells and (B) proportion of FOXP3+ $T_{reg}$ cells in the mesenteric lymph nodes (MLNs) harvested from *B. fragilis* WT or Δ*nanH* mono-colonized mice. Cells were collected 4 weeks after mono-colonization, stained, and analyzed by flow cytometry. n=10 per group. Two-way ANOVA: **P*<0.05

**Supplemental Tables**

Table 1. Proteomics of *B. fragilis* in pooled HMOs.

Table 2. NanH operon in *B. fraglis* strains.

Table 3. Bacterial strains used in this study.

Table 4. Primers used in this study.

**Table 3**. Strains and plasmids used in the study.

| Bacterial Species | Strain | Description | Reference |
|---|---|---|---|
| *B. fragilis* | NCTC 9343 | Type strain | 1 |
| *B. thetaiotaomicron* | VPI 5482 | Type strain | 2 |
| *B. vulgatus* | ATCC 8482 | Type strain | 3 |
| *B. ovatus* | ATCC 8483 | Type strain | 3 |
| *B. salyersiae* | DSM 18765 | Type strain | 4 |
| *B. uniformis* | ATCC 8492 | Type strain | 3 |
| *B. acidifaciens* | JCM 10556 | Type strain | 5 |
| *B. fragilis* ΔCCF | | *B. fragilis* lacking bf3579-bf3583 | 6 |
| *B. fragilis* Δ*nanH* | | *B. fragilis* lacking bf1806 | This study |
| *B. fragilis* ΔCCFΔ*nanH* | | *B. fragilis* lacking bf3579-bf3583 and bf1806 | This study |
| *E. coli* CC118 lambda pir | *Δ(ara-leu) araD ΔlacX74 galE galK phoA20 thi-1 rpsE rpoB argE (Am) recA1 λpir* | Plasmid propagation | 7 |
| *E. coli* S-17 lambda pir | *recA pro hsdR RP4- 2 (Tc::Mu;Km::Tn7) ( λ pir)* | Conjugal transfer | 7 |
| PKNOCK-bla-erm | | Bacteroides suicide vector, mob+, Tra-, AmpR (*E. coli*), ErmR (*Bacteroides*) | 8 |
| pFD340 | | Bacteroides shuttle vector, contains IS4351 promoter, AmpR (*E. coli*), ErmR (Bacteroides) | 9 |
| pFD340-cat | | Modified pFD340 plasmid, AmpR (E. coli), CmR ErmR (*Bacteroides*) | 6 |
| pFD340-tetQ | | Modified pFD340 plasmid, AmpR (*E. coli*), TetR ErmR (*Bacteroides*) | 6 |

**Table 4. Primers used in the study.**

| Primer target | | Primer sequence (5' > 3') | Reference |
|---|---|---|---|
| ***Primers for quantitative PCR*** | | | |
| *B. fragilis* NCTC 9343 | bf2779 | Forward GTACACTGCTCGAGATTATG<br>Reverse GTCGTCCTGAAACACATAG | This study |
| *B. thetaiotaomiron VPI-8482* | bt3780 | Forward CATCGAAGGGATGGTTTATT<br>Reverse CAGGTTTCTTCGGATTGTAG | This study |
| *B. vulgatus* ATCC 8482 | 16S | Forward TCATCGTGGTCCATTGTCGG<br>Reverse AACACCCCGTCAAAATTGCG | This study |
| *B. fragilis* ccfC | bf3851 | Forward GATGAACTGATAGCCCATTA<br>Reverse TAGCGATGACTAAAGGTGTT | 1 |
| *B. fragilis* ccfD | bf3580 | Forward CCAGTT CCG TCC CTC TAT TATT<br>Reverse ACTCTC GCGTCA TTA GGA TTG | This study |
| *B. fragilis* ccfE | bf3579 | Forward TGCTATTTGCACGGGTAACA<br>Reverse CCGAAACTCCGA TTCTTCAT | 1 |
| *B. fragilis* gyrB | gyrase B | Forward GTGAATGAGGACGGCAGTTT<br>Reverse CTCGATGGGGATGTTTTGTT | 2 |
| *B. fragilis nanH_2* | bf4051 | Forward ATACGGCCCAGTTGGTATTG<br>Reverse GGCTCTTTCACCTGATCTGTT A | This study |
| *B. fragilis* BF3412 | bf3412 | Forward GCACCGAGCTGGGATATAAA<br>Reverse ACACTCCGACAGAAGGATAGA | This study |
| *B. fragilis* BF3413 | bf3413 | Forward GTCCAAACCCTGGACGATTAC<br>Reverse GGCAAAGCGCATCCATTTATT | This study |
| *B. fragilis* BF3414 | bf3414 | Forward ACTTCTTGCTCACGCGATTA<br>Reverse GTGGATAGAGCCGTTCAGATG | This study |
| *B. fragilis* BF1802 | bf1802 | Forward GGACCTTTATGGCCTCATACA<br>Reverse GCTCCCTGTGCATTTACATAAC | This study |
| *B. fragilis* BF1806 | bf1806 | Forward TGAGTGACGGTACTTTGGTATTC<br>Reverse AGTTCTTGCCACCATCTTTACT | This study |
| *B. fragilis* BF1807 | bf1807 | Forward CATTGGTGCGGGTATCTATCTT<br>Reverse CAGCCCTACTTCCTGATCTTTC | This study |
| *B. fragilis ΔnanH* | *ΔnanH* | Forward ACTATCTTTGCTCCCGATAACA<br>Reverse CCGGGAGCAGGTGATATTC | This study |
| Universal 16S | 16S | Forward ACTCCTACGGGAGGCAGCAGT<br>Reverse ATTACCGCGGCTGCTGGC | 1 |

| Primer target | | Primer sequence (5' > 3') | Reference |
|---|---|---|---|
| ***Primers for generating deletion constructs*** | | | |
| Δbf1806 Flank 1 | 1 kb region upstream of bf1806 | Forward ACCGCGGTGGCGGCCGCTCTAGAACTAG TGGGCACAGAATGTATGGGG Reverse CCTGCCCGAACCGAAGGTATCGACTAATT AATATTTATGTTATCG | This study |
| Δbf1806 Flank 2 | 1 kb region downstream of bf1806 | Forward TAATTAGTCGATACCTTCGGTTCGGGCAG GTGAATATC Reverse AGCTTGATATCGAATTCCTGCAGCCCGG GGATCAATCTCGCCAGGTTTATAATGC | This study |
